# Benchmarking feature selection and feature extraction methods to improve the performances of machine-learning algorithms for patient classification using metabolomics biomedical data

**DOI:** 10.1101/2023.12.21.572852

**Authors:** Justine Labory, Evariste Njomgue-Fotso, Silvia Bottini

## Abstract

**Objective:** Classification tasks are an open challenge in the field of biomedicine. While several machine-learning techniques exist to accomplish this objective, several peculiarities associated with biomedical data, especially when it comes to omics measurements, prevent their use or good performance achievements. Omics approaches aim to understand a complex biological system through systematic analysis of its content at the molecular level. On the other hand, omics data are heterogeneous, sparse and affected by the classical “curse of dimensionality” problem, i.e. having much fewer observation samples (*n*) than omics features (*p*). Furthermore, a major problem with multi-omics data is the imbalance either at the class or feature level. The objective of this work is to study whether feature extraction and/or feature selection techniques can improve the performances of classification machine-learning algorithms on omics measurements.

**Methods:** Among all omics, metabolomics has emerged as a powerful tool in cancer research, facilitating a deeper understanding of the complex metabolic landscape associated with tumorigenesis and tumor progression. Thus, we selected three publicly available metabolomics datasets, and we applied several feature extraction techniques both linear and non-linear, coupled or not with feature selection methods, and evaluated the performances regarding patient classification in the different configurations for the three datasets.

**Results:** We provide general workflow and guidelines on when to use those techniques depending on the characteristics of the data available. For the three datasets, we showed that applying feature selection based on biological previous knowledge improves the performances of the classifiers. Notebook used to perform all analysis are available at: https://github.com/Plant-Net/Metabolomic_project/.

## Introduction

Personalized medicine concerns the development of approaches able to stratify patients based on their disease subtype, risk, prognosis, or treatment response using specialized diagnostic tests [1]. The key idea is to identify medical decision elements based on individual patient characteristics, including molecular biomarkers, rather than on population averages [2]. Lately, the development of precision medicine has seen unprecedented growth, thanks to the development of omics technologies and machine learning approaches [3].

Omics technologies provide a global view of the molecules that compose a cell, a tissue or an organism. They are mainly aimed at the universal detection of genes (genomics), mRNAs (transcriptomics), proteins (proteomics) and metabolites (metabolomics) in a specific biological sample [4]. The fundamental aspect of these approaches is that a complex system can be understood more thoroughly if it is considered as a whole. Each omics represents a layer of information of this complex system and the objective is to study the biological mechanisms in their entirety and the complexity of their interactions.

Modern metabolomics produces high-dimensional datasets comprising hundreds or even thousands of measured metabolites in large-scale human studies involving thousands of participants [5]. One of the key goals of metabolomics, mainly when applied in cancer research, is the discovery of robust and reliable biomarkers for early detection, diagnosis, prognosis, and treatment response prediction. Traditionally, cancer biomarker discovery has focused on genomic and proteomic approaches; however, metabolomics offers several advantages in this regard [6]. Metabolites represent the downstream products of cellular processes, capturing the integrated effects of genetic and environmental factors, as well as dynamic changes in the tumor microenvironment [7]. Moreover, metabolites are accessible through minimally invasive techniques, such as blood or urine sampling, enabling their potential translation into clinical practice [8].

The drawback that prevents wider use of metabolomics, as well as for other omics, is that data collection is financially costly, and the number of clinical research participants is usually limited. This yields unbalanced datasets in which the number of metabolites measured (features) far exceeds the number of patients (observations) [9]. This issue is known as the curse of dimensionality. Also, with many features, learning models tend to overfit, which may cause performance degradation on unseen data. Furthermore, most of the features are highly correlated and some features are not always directly connected with disease explanation, thus resulting in a high-dimensional space composed of many redundant and non-informative features that can mislead the algorithm training. Therefore, extracting systemic effects from high-dimensional datasets requires dimensionality reduction approaches to untangle the high number of metabolites into the processes in which they participate.

Dimensionality reduction is one of the most powerful tools to address the previously described issues. It can be mainly categorized into two main components: feature selection and feature extraction. Feature selection finds a subset of the *original* features that maximise the accuracy of a predictive model [10]. It can be based on prior knowledge such as evidence from known literature or based on existing databases [11,12]. Feature extraction methods project the original high-dimensional features to a new feature space with low dimensionality. The newly constructed feature space is usually a linear or nonlinear combination of the original features. Among the different techniques of feature extraction, we focused here on latent representation learning, which is a machine learning technique that attempts to infer latent variables from empirical measurements [13]. Latent components also called latent space, in contrast to observed variables, are information that is not measurable therefore have to be inferred from the empirical measurements. Several techniques have been developed to infer the latent space with successful applications on omics data however, how to choose the model that fits the best with the available data is very challenging.

While during recent years there has been a lot of enthusiasm about the potential of ‘big data’ and machine learning-based solutions, there exist only a few examples that impact current clinical practice [14]. The lack of impact on clinical practice can largely be attributed to insufficient performance of predictive models, difficulties to interpret complex model predictions, and lack of validation via prospective clinical trials that demonstrate a clear benefit compared to the standard of care. The objective of our work is to explore the performances in patient classification based on their metabolomics profile of several linear and non-linear techniques of feature extraction and feature selection and to provide general guidelines on when to use those techniques depending on the data available.

## Related works

Latent space representations in metabolomics have been applied to classify individuals based on their phenotype (i.e. healthy/disease). Nyamundanda et al. successfully used Probabilistic Principal Component Analysis (PPCA) to identify metabolites which were responsive to pentylenetetrazole (the treatment used in the study) [15]. They also used a mixture of PPCA (MP- PCA) to simultaneously cluster and reduce the dimension of metabolites data. They have demonstrated that the application of those techniques helps in the identification of disease phenotypes or treatment-responsive phenotypes. Gomari et al. trained a VAE model on 217 metabolite measurements in 4644 blood samples from the TwinsUK study [16]. They analysed the features’ importance with Shapley Additive Global Importance (SAGE) technique at different levels such as metabolites, sub-pathways, and super-pathways. They showed that VAE latent dimensions capture a complex mix of functions related to sub-pathways, thus capturing major metabolic processes in the dataset. Chardin et al. [17] proposed a supervised Auto Encoder (SAE) which provided an accurate localization of the patients in the latent space, and an efficient confidence score. The confidence score is the probability of the diagnosis, which gives additional information to the clinician about the classification. The SAE also included a feature selection step, which was able to identify the metabolites known to be biologically relevant.

Such approaches have been applied also in metabolomics studies beyond human pathologies. Hamzehzarghani et al. [18] used factor analysis to profile the metabolic of spikelets of wheat cultivars, Roblin and Sumai3, susceptible and resistant to fusarium head blight, respectively. By combining factors and factors loading they were able to identify metabolites involved in pathogen-stress and their metabolic pathways of synthesis. Date et al. [19] described an improved DNN-based analytical approach that incorporated an importance estimation for each variable using a mean decrease accuracy (MDA) calculation. The performance of the DNN-MDA approach was evaluated using a data set of metabolic profiles derived from yellowfin goby that lived in various rivers throughout Japan.

## Materials and Methods Datasets

We have used three metabolomics datasets from three different types of cancer: brain, breast, and lung cancers, whose characteristics are summarized in Table 1.

**Table 1:** Description of the characteristics of the metabolomic datasets used in this study. A comprehensive table summarizing key features, such as the number of metabolites and samples, the different sample types and classes in three cancer datasets.

| Dataset | # Metabolites | # Samples | Type of sample | Classes | References |
| --- | --- | --- | --- | --- | --- |
| BRAIN | 7017 | 88 | Glial tumor tissue | IDH wild-type tumor/<br>IDH-mutant tumor | Chardin *et al.* BMC Bioinformatics (2022) |
| BREAST | 162 | 271 | Tumor tissue | ER+ tumor/ER- tumor | Budczies J *et al.* J Proteom. (2013) |
| LUNG | 2944 | 1005 | Urine | Cancer patient/<br>Healthy patient | Mathé E, et al. Cancer Res. (2014) |

### BRAIN dataset

The BRAIN dataset consists of 7017 metabolites from 88 samples of glial tumors: 38 isocitrate dehydrogenase (IDH) wild-type tumors and 50 IDH-mutant tumors. Tumor samples were analyzed in an unbiased metabolomics using Liquid Chromatography coupled to tandem Mass Spectrometry (LC-MS/MS) [17].

### BREAST dataset

The BREAST dataset includes 162 metabolites from 271 breast cancer tissues: 204 samples which have receptors for estrogen (ER+) and 67 samples which do not have receptors for estrogen (ER−) [20]. The metabolomic analysis was performed by gas chromatography followed by time-of-flight mass spectrometry (GC–TOFMS) as described here [21].

### LUNG dataset

The LUNG dataset is composed of 2944 metabolites concerning urine samples from 469 Non-Small Cell Lung Cancer (NSCLC) patients prior to treatment and 536 controls [22]. The dataset was obtained after an unbiased liquid chromatography/mass spectrometry approach. It is available at MetaboLights (study identifier MTBLS28).

It is important to notice the very different characteristics of the three metabolomic datasets, both in terms of the number of features and number of patients (Table 1). The BRAIN dataset contains a limited number of patients (88) with several features (7,017). The BREAST dataset contains a moderate number of patients (271) with a small number of features (162). The LUNG dataset contains a very large number of patients (1,005) with a large number of features (2,944).

### Feature selection method

We define feature selection here as a data preprocessing strategy aimed at finding a subset of the original features that maximize the accuracy of a predictive model. The aim is to prepare understandable, clean data in order to build a simpler, more comprehensible model. It can be based on prior knowledge, i.e., evidence from known literature, or biological properties of the studied system. We have used the Kolmogorov–Smirnov test to identify the features with the most significant difference between the two populations (e.g., diseased vs healthy).

To apply the KS test, we used the function “ks.test” from the “dgof” R package [23].

### Feature extraction methods

We tested linear and non-linear techniques. Linear techniques suppose that there is a linear relationship between the observed variables and the latent space. Under this assumption, the latent space can then be inferred from observed variables. We test four linear methods: Principal Component Analysis (PCA), Mixture of Probalistic PCA (MPPCA), High dimensional discriminant analysis (HDDA) and Factor Analysis (FA) and two non-linear: Kernel PCA (KPCA) and Gaussian Process Latent Variable Modeling (GPLVM). We have also run these latent space inference methods on pre-selected features to analyze the impact of this preprocessing on the performance metrics.

### Linear techniques

#### Principal Component Analysis (PCA)

PCA helps to identify the most important patterns or trends in the data and represents them using a smaller number of new variables called principal components. First, the data are standardized to ensure all variables have a similar scale. Then, the covariance matrix is computed which measures how the variables are related to each other. Then the eigenvalues and eigenvectors of the covariance matrix are also calculated. Eigenvalues represent the amount of variance explained by each eigenvector, while eigenvectors represent the direction or pattern in the data, also called principal components. Each principal component represents a linear combination of the original variables, weighted by the corresponding eigenvector.

#### Mixture of probalistic PCA (MPPCA)

MPPCA is a probabilistic model that combines multiple probabilistic PCA (PPCA) models into a mixture model. MPPCA extends the concept of PPCA to capture more complex data distributions and capture data points that may belong to different clusters or components. PPCA is a linear dimensionality reduction technique that assumes a linear relationship between the observed variables and a lower-dimensional latent space. PPCA assumes that the observed data points are generated by adding Gaussian noise to a low-dimensional subspace, which is represented by a linear mapping from the latent space to the observed space. In MPPCA, a mixture model framework is employed to account for multiple components in the data. MPPCA assumes that each observed data point is associated with a latent variable, which indicates the component to which it belongs. The model further assumes that the latent variables follow a certain probability distribution, such as a Gaussian distribution. The mixing coefficients represent the probability of a data point belonging to each component.

#### High dimensional discriminant analysis (HDDA)

HDDA extends traditional Linear Discriminant Analysis (LDA) to handle situations where the number of variables or features is large compared to the number of samples. LDA is a classical technique that aims to find a linear transformation of the data that maximizes the separation between different classes or groups. LDA assumes that the data follows a multivariate Gaussian distribution with class-specific means and a shared covariance matrix. HDDA addresses the challenges of high-dimensional data by incorporating regularization and shrinkage techniques. Regularization methods are employed to stabilize the estimation of the covariance matrix. HDDA performs dimensionality reduction by projecting the high-dimensional data onto a lower-dimensional subspace. This subspace is determined by a set of linear discriminant directions that maximize the separation between classes. The number of discriminant directions is typically smaller than the original dimensionality, allowing for a reduced representation of the data. After dimensionality reduction, HDDA can be used for classification tasks. New samples can be projected onto the reduced subspace, and their class labels can be predicted based on their proximity to the class-specific centroids or by using other classification algorithms.

#### Factor Analysis (FA)

FA is a statistical method that analyzes the relationships among observed variables to uncover the latent factors that explain their covariation. It assumes that the observed variables are influenced by a smaller number of unobserved factors, also known as common factors. First, the correlation matrix from the observed variables is calculated. This matrix represents the pairwise relationships and covariation among the variables. Then, the factor extraction step identifies the underlying factors that explain the observed covariation. However, the application of FA requires certain conditions: the observed variables must be highly correlated and the number of samples must be at least four times greater than the number of features.

To check that if the variables in our datasets are highly correlated, we run the Kaiser-Mayer-Olkin (KMO) test with the function KMO from the R package “EFAtools” [24]. If the KMO value is higher than 0.7, then FA can be performed, otherwise, it is not possible.

Then to define the number of factors to set for FA, we use the function “N_FACTORS” from “EFAtools” package which allows us to perform various factor retention criteria simultaneously. We selected the most frequently used number of factors.

#### Non-linear techniques

Non-linear techniques assume that the relationship between the latent space and observed variables is not linear. However, some non-linear techniques can, under some constraints, help to infer linear relationship while linear techniques can only infer linear relationships. We tested 2 non-linear methods: Kernel PCA (KPCA) and Gaussian Process Latent Variable Modeling (GPLVM).

#### Kernel PCA (KPCA)

KPCA is a non-linear extension of PCA. It allows for capturing non-linear relationships and patterns in high-dimensional data. KPCA operates by mapping the data into a higher-dimensional feature space using a kernel function and then performing linear PCA in that space. KPCA begins by selecting an appropriate kernel function which calculates the similarity or distance between data points in the original input space. Each data point is then transformed or mapped into a higher-dimensional feature space using the kernel function. In this higher-dimensional feature space, the covariance matrix is calculated based on the mapped data points. This covariance matrix represents the relationships and variances among the transformed data. The data are projected onto the principal components, which are the eigenvectors of the covariance matrix. The projection yields a lower-dimensional representation of the data, where the dimensions correspond to the principal components. Finally, the principal components that capture the most significant patterns and variance in the data are selected.

#### Gaussian Process Latent Variable Modeling (GPLVM)

GPLVM is a probabilistic dimensionality reduction technique that combines Gaussian processes with latent variable models. GPLVM aims to learn a low-dimensional representation of high-dimensional data by modeling the underlying structure and uncertainty in the data.

GPLVM assumes that the observed high-dimensional data points are generated from a lower-dimensional latent space. Each data point is associated with a set of latent variables that lie in the lower-dimensional space. Gaussian processes are non-parametric models that can represent complex functions. In GPLVM, a Gaussian process is used to model the mapping from the latent space to the observed space. This mapping represents how the latent variables influence the observed data. Then, GPLVM employs Bayesian inference to estimate the latent variables and the parameters of the Gaussian process. It aims to find the most likely values of the latent variables given the observed data. Once the latent variables are estimated, GPLVM provides a reduced-dimensional representation of the data. This lower-dimensional representation retains the most important information and captures the underlying structure in the data.

All the linear and non-linear techniques were implemented in R and only XGBoost classification were implemented in python. The only difference is for methods which have its own classification namely GPLVM, HHDA and MPPCA, a 4-fold cross validation approach was used and for other methods with XGBoost classification, it was a five repeated 4-fold cross validation approach.

### Classification

For feature extraction techniques that do not perform classification already in their model, we used the XGBoost classifier model [25]. XGBoost is an implementation of gradient-boosting decision trees sequentially combine decision trees to create an ensemble model, particularly used for classification and regression. It was selected for its speed, scalability, and superior performance, making it a popular choice in various analyses.

Due to the different inherent characteristics of datasets, we have defined two workflows for classification with XGBoost explained hereafter.

#### LUNG workflow

We divided the original dataset in two partitioned datasets: 70% of the data for training and 30% of the data for testing. Then, we used a 5-fold cross-validation approach on the training dataset to tune the parameters of XGBoost and used the test dataset to assess the predictions.

#### BREAST and BRAIN workflow

BREAST and BRAIN datasets are unbalanced datasets. They are characterized by a disproportionate distribution of instances between different classes, and they are often an obstacle for traditional machine learning models, as they tend to adopt a behavior biased in favor of the majority class. To overcome this limitation and ensure the development of a predictive model capable of discerning patterns in unbalanced data, a repeated 4-fold cross-validation approach was employed with default parameters of XGBoost. This technique involves partitioning the dataset into four subsets, utilizing three of them for training and the remaining one for validation in each iteration. This process is repeated four times, with each fold serving as the validation set in a distinct iteration. To ensure a more robust evaluation of the model’s performances, we repeated this process 5 times, it provides multiple independent estimates of how well the model generalizes to unseen data. Finally, the model undergoes training and evaluation several times, enabling a more thorough assessment of its predictive capabilities in various data subsets. By applying this strategy, we aimed to mitigate the impact of unbalanced class distribution, ensuring that the XGBoost model learns from patterns effectively and generalizes well to novel and unseen instances, thus contributing to the reliability and efficiency of the predictive modeling process.

### Calculation of metrics for performances evaluation

We have chosen to report seven metrics to evaluate performances of the models.

The accuracy is the number of correct predictions whether positive or negative, defined as:

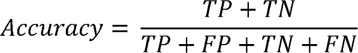

Where True Positive (TP) is the number of correct positive predictions, False Positive (FP) is the number of incorrect positive predictions, True Negative (TN) is the number of correct negative predictions and False Negative (FN) is the number of incorrect negative predictions.

The precision quantifies the number of correct positive predictions out of the positive predictions made by the model:

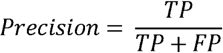

The recall, also called the sensitivity, is the number of TP among the real positive samples (TP and FN) that the model obtains, calculated with the following formula:

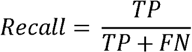

The specificity is the number of correct negative predictions the model can detect.

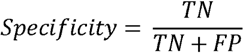

The F1 score keeps the balance between precision and recall. It’s often used when class distribution is uneven.

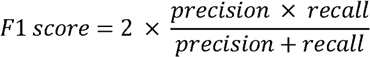

The AUC measures the performance of a binary classification model by quantifying the Area Under the Receiver Operating Characteristic (ROC) curve.

For the two unbalanced datasets we calculated the balanced accuracy. It’s the arithmetic mean of sensitivity and specificity.

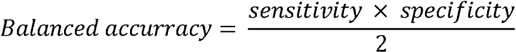

For all these metrics, we have computed a 95% confidence interval. For LUNG dataset, we used the boostrap method with 20 iterations. For BREAST and BRAIN datasets, we computed the confidence interval on results of the 4-folds repeated five times of the cross validation.

## Computational workflow

We set up a workflow consisting of 2 main steps and an optional step (figure 1). The two main steps are first the execution of a feature extraction model and then the classification of patients using the newly calculated features.We also performed feature selection (optional step) before the feature extraction model and run again the analysis.

**Figure 1:**
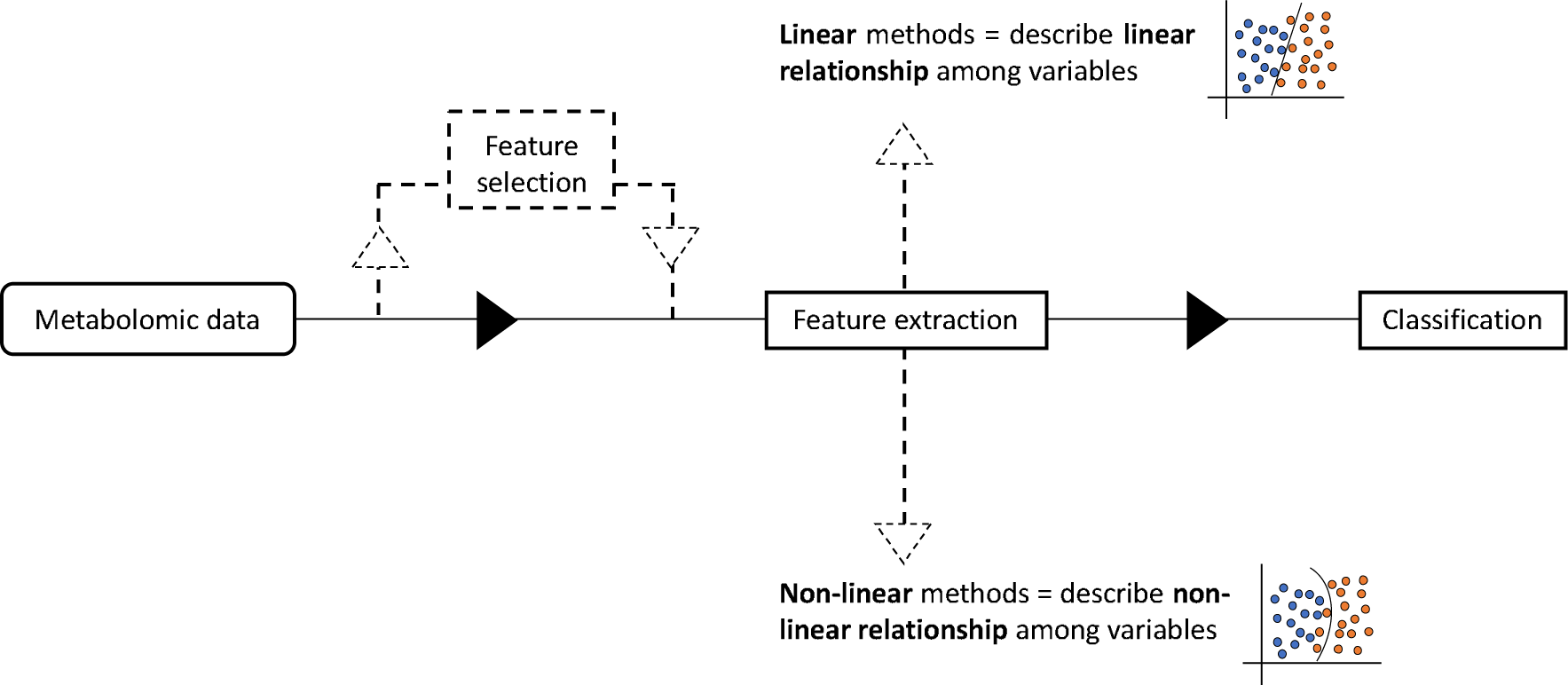
Workflow used in this study to evaluate the classification performances of several feature extraction techniques coupled or not with feature selection. The input of the workflow consists in the metabolomics dataset, which is a table containing the concentrations of different metabolites in each sample of each dataset. The workflow consists of 3 steps: one optional and 2 mandatories. The first step is the feature selection, which is optional that consists in selecting some of the original features based on the Kolmogorov–Smirnov test to identify the features with the most significant difference between the two classes of samples. The second step is the feature extraction. We included four linear and two non-linear methods. Finally, for sample classification, we used the XGBoost model for methods that do not have their own classification method.

Then the performances on the classification task are evaluated using the aforementioned metrics. When possible, we also calculated the features importance to see whether the model identify the correct biomarkers in each dataset (only for PCA, FA, and XGBoost).

To compute feature importance, we used SHAP (Shapley Additive exPlanations) values [26]. SHAP values allow us to attribute a specific contribution to each feature for a given prediction and to understand the unique role each feature plays in influencing the model’s decisions. By leveraging SHAP values, we gain a comprehensive and interpretable view of feature importance, contributing to a more informed and transparent analysis of the XGBoost model’s predictive capabilities.

Overall, we trained four linear and two non-linear models by feeding with either all the measured metabolites as features or after performing feature selection. These models were compared to the reference model called NFE for No Featured Extraction that consists in using only XGBoost classifier with or without feature selection.

Notebook used to perform all analysis are available at: https://github.com/Plant-Net/Metabolomic_project/

## Results

We applied the experimentation workflow to the three metabolomic datasets as described in the methods section and in Figure 1. The main characteristics of the datasets are reported in Table 1. Briefly, the BRAIN dataset contains a limited number of patients (88) with high number of metabolites (features) (7,017). The BREAST dataset contains a moderate number of patients (271) with a small number of features (162). The LUNG dataset contains a very large number of patients (1,005) with a large number of features (2,944). When feature selection is applied before feature extraction, a smaller number of metabolites is used, namely: 269 metabolites for the BRAIN dataset, 52 metabolites for the BREAST dataset and 202 metabolites for the LUNG dataset.

### Cross-validation helps to handle unbalanced and small datasets

When it comes to overcoming the challenges posed by small and unbalanced datasets, cross-validation emerges as a crucial and indispensable strategy in machine learning. In scenarios with unbalanced class distributions or limited data, traditional model evaluation can lead to biased and unreliable results. Cross-validation, with its ability to iteratively partition the dataset into training and test subsets, offers a more robust solution. By ensuring that each data point participates several times in the evaluation process, cross-validation provides a more complete understanding of a model’s performance. This approach is particularly valuable when dealing with unbalanced datasets, where instances of minority classes may be overlooked. Furthermore, in the context of small datasets, cross-validation maximizes the usefulness of limited samples by systematically evaluating model performance in different partitions. In our study we showed the importance of using cross-validation on small, unbalanced datasets. The BRAIN dataset is the smallest dataset in our analysis and is unbalanced. By using cross-validation, we achieved good performances with a maximum of F1 score of 90.4 (±6.5) (supplementary table 1). Although the BREAST dataset contains more samples than the BRAIN dataset, it is very unbalanced. With cross-validation, we were able to achieve a F1 score of 94.0 (±2.7) (supplementary table 2).

Finally, cross-validation is a powerful tool for addressing the complex problems posed by small, unbalanced datasets, contributing to the development of more reliable and generalizable machine learning models.

### Feature selection improves classification performances on all datasets

To evaluate the performances of the different model, we used the accuracy or balanced accuracy for unbalanced datasets, the recall, the specificity and the F1 score because together they give an overview of model’s performances. A high precision can mask poor performances in capturing positive instances corresponding to low recall, thus a focus on F1 score helps us to find a balance between precision and recall, while specificity gives us an idea of the rate of true negatives.

In supplementary table 1, we summarize all the results for the BRAIN dataset. We could not perform FA because the KMO value was equal to 0.56, thus smaller than the required threshold for applicability. The best performances were obtained with No Feature Extraction method (NFE) coupled with feature selection achieving an average balanced accuracy of 89.6% (±6.9%), an average recall of 92.1% (±7.8%), an average specificity of 88.9% (±12.0%) and an average F1 score of 90.4% (±6.5%) (figure 2A). The feature selection before applying XGBoost classification slightly improved the performances, suggesting that XGBoost already captures the important features that drive the classification. It is interesting to note that the non-linear GPLVM method achieved the best performances among the feature extraction methods with an average balanced accuracy of 87.7% (±8.4%), an average recall of 86.2% (±16.9%), an average specificity of 89.2% (±9.1%) and an average F1 score of 88.3% (±10.4%). The GPLVM performances are very similar to NFE performances, but the standard deviations are bigger, meaning that the GPLVM model has greater variability. Although we observe that feature selection improves the performances of all feature extraction methods, KPCA obtained very poor scores, almost comparable to a random classifier.

**Figure 2:**
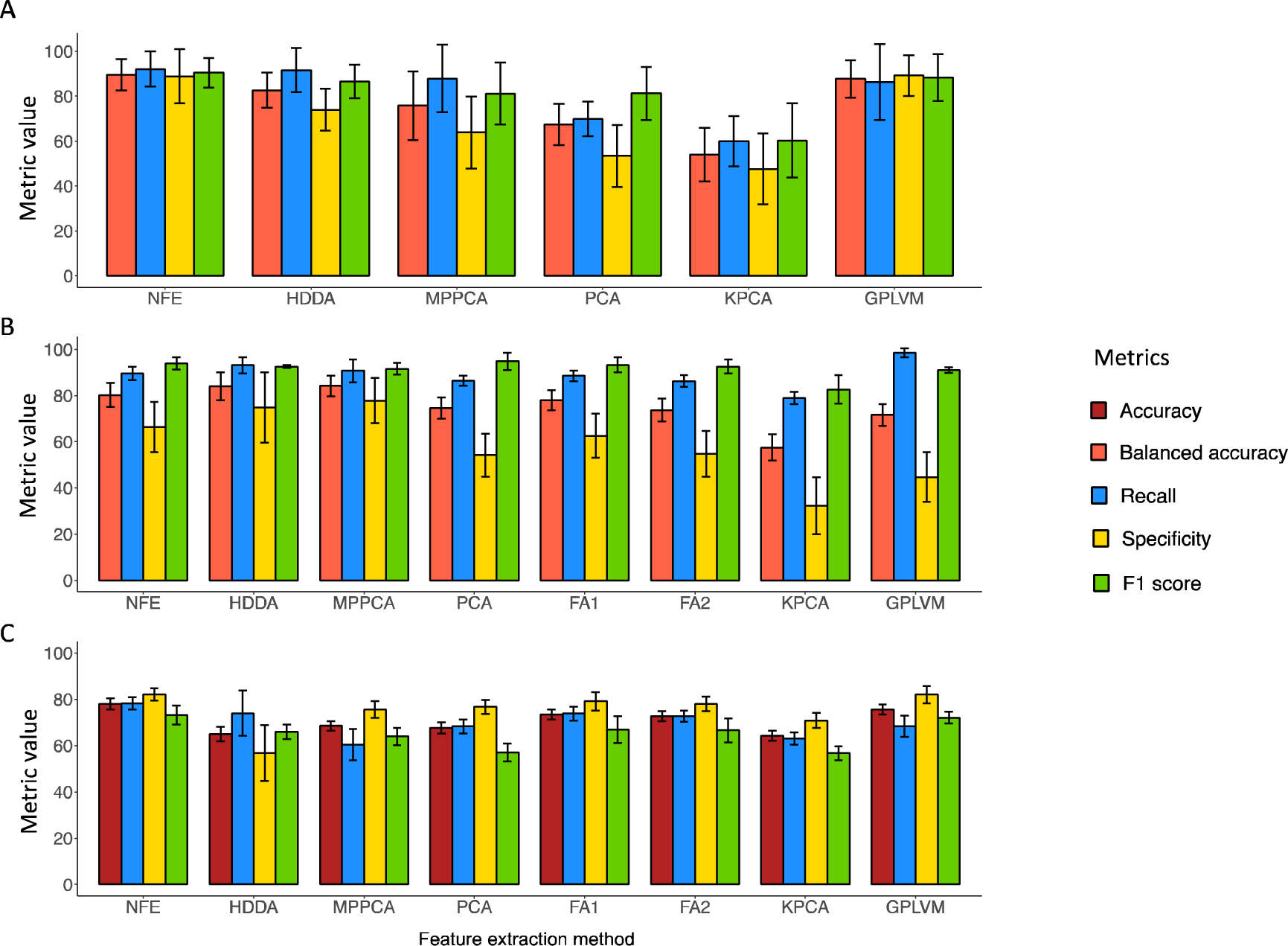
Metrics to evaluate the performances of the model used in this study. Performances after the feature selection step of each extraction method in terms of accuracy (dark red), balanced accuracy (salmon), recall (blue), specificity (yellow) and F1 score (green) for BRAIN dataset (A), BREAST dataset (B) and LUNG dataset (C). The abbreviations for feature extraction methods are: NFE: No Feature Extraction; HDDA: High dimensional discriminant analysis; MPPCA: Mixture of Probalistic PCA; PCA: Principal Component Analysis (PCA); FA: Factor Analysis; KPCA: Kernel PCA; GPLVM: Gaussian Process Latent Variable Modeling. For BRAIN and BREAST dataset, two FA were performed with different numbers of features. For BREAST dataset, seven feature were used for FA1 and ten for FA2. For LUNG dataset, 17 features were used for FA1 and 44 for FA2. For BRAIN, we were unable to perform FA.

Regarding the BREST dataset, the best performances are achieved by combining feature selection and the linear method HDDA with an average balanced accuracy of 84.0% (±6.0%), an average recall of 93.1% (±3.4%), an average specificity of 74.8% (±15.3%) and an average F1 score of 92.5% (±0.6%) (figure 2B and supplementary table 2). We can also observe that similar scores were obtained with MPPCA after feature selection. Thus, the application of a feature extraction method before sample classification improves performances. Comparably to the results on the previous dataset, KPCA yielded the worst performances although the overall scores are better than the ones obtained on the previous dataset.

Finally, for the LUNG dataset, the method that achieved the best performances is the combination of feature selection and XGBoost model as for the BRAIN dataset, with an average accuracy of 78.4% (±2.4%), an average recall of 78.7% (±2.9%), an average specificity of 82.5% (±2.6%) and an average F1 score of 73.8% (±3.1%) (refer to supplementary table 3 for results for the LUNG dataset). For this dataset, achieving a high recall rather than high precision is crucial as we seek to distinguish healthy from cancer individuals, hence patients erroneously predicted as healthy (FNs) can have dramatic consequences. Among feature extraction methods, the non-linear GPLVM model perform the best with an average accuracy of 75.6% (±1.8%), an average recall of 67.2% (±2.9%), an average specificity of 83.0% (±3.2%) and an average F1 score of 71.9% (±2.2%) (figure 2C). As for the other datasets, performing feature selection before feature extraction improved performances, independently by the employment of a feature extraction model.

In summary, for two datasets out of three, the combination of feature selection and NFE method achieves the best performances. While for BRAIN and LUNG datasets feature selection combined with classification achieved the best performances, for BREAST dataset, the combination of feature selection and the HDDA extraction method yields the best achievement. Of note, the latter dataset is the only dataset for which the number of samples exceeds the number of features that might point out the need for feature extraction combined to feature selection. Overall, we can observe that feature selection before feature extraction always improve classification metrics.

### ROC curves as a useful tool to compare model performances among multiple datasets

The ROC curve and its associated AUC serve as critical tools for evaluating and comparing the performance of different classification methods. The AUC ROC curve evaluates the ratio between a model’s true-positive and false-positive rates for different threshold values, providing an overview of its discriminatory power. A higher AUC value indicates better model performance, as it means a greater ability to distinguish positive from negative instances.

Since the feature selection significantly improve classification performances, we calculated ROC curves for all feature extraction methods and NFE model after feature selection. For the BRAIN dataset, performances varie widely from one method to another (figure 3A). The two methods that best perform are NFE and GPLVM. The worst-performing method is KPCA, whose performances are close to a random classifier for this dataset. Regarding the BREAST dataset, unlike the BRAIN dataset, all the methods, except for KPCA, achieved comparable results (figure 3B). For the LUNG dataset, all methods achieved comparable performance (Figure 3C).

**Figure 3:**
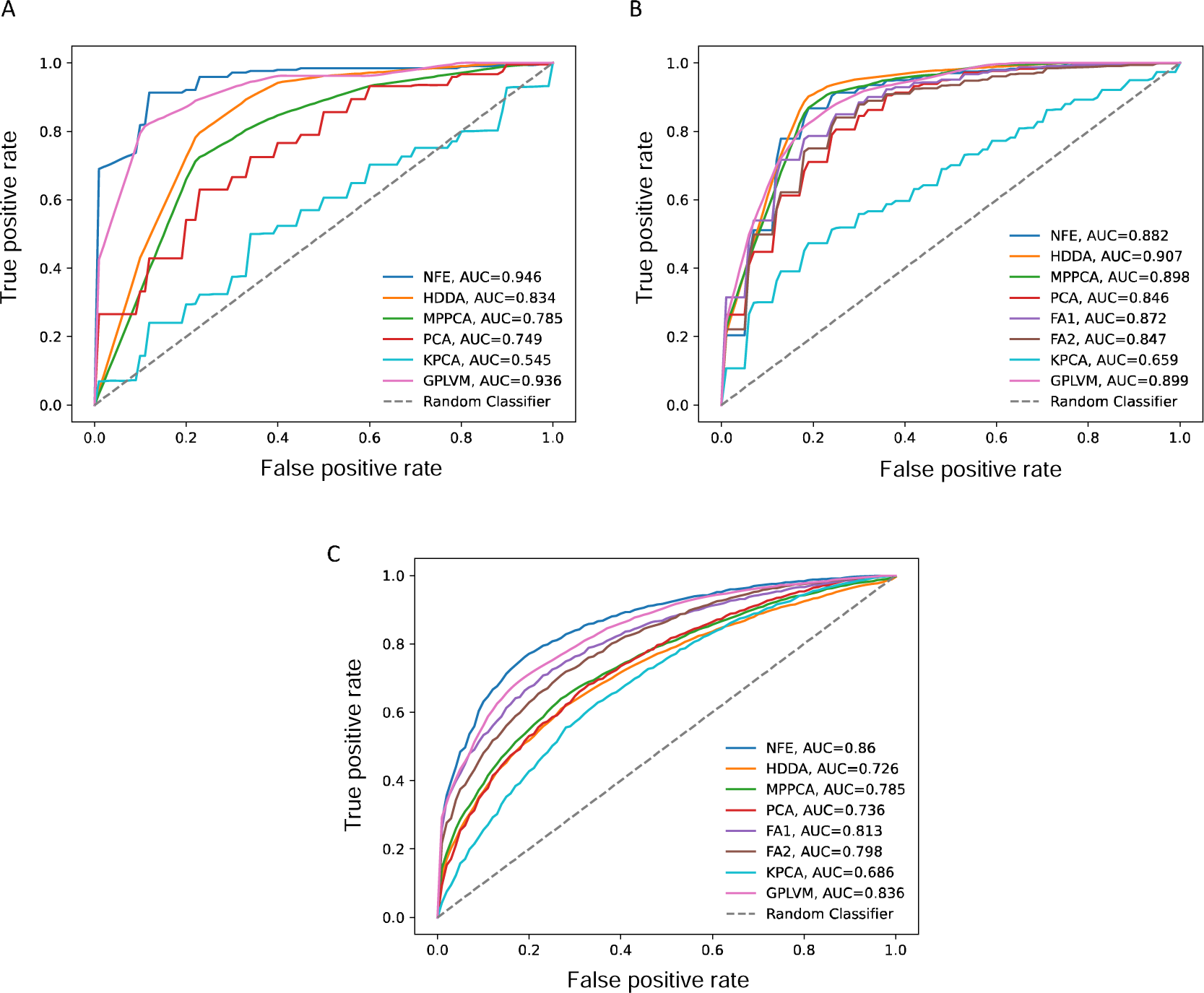
ROC curves to compare multiple model performances on multiple datasets. ROC curves obtained by performing feature selection and feature extraction are shown for BRAIN dataset (A), BREAST dataset (B) and LUNG dataset (C). The blue curve is performances of no feature extraction (NFE) method, the orange one is for High dimensional discriminant analysis (HDDA), the green one for Mixture of Probalistic PCA (MPPCA), the red one for Principal Component Analysis (PCA), the purple one and brown one for Factor Analysis (FA), the cyan one for Kernel PCA (KPCA) and the pink one for Gaussian Process Latent Variable Modeling (GPLVM). The dashed grey line corresponds to performances of a random classifier.

In summary, the KPCA method is the worst performer, suggesting that is not suitable for extracting relevant features in the context of metabolomics cancer data. There is no clear difference in performances between linear and non-linear methods. The most stable method, independently on the dataset, appears to be NFE, namely the use of XGBoost model coupled with feature selection to classify samples.

### Features importance allows to identify potential biomarkers

As last step of our analysis we inspected features importance of the best performing models for each dataset using SHAP explainer [26]. By calculating the features contribution to the classification, we can identify the metabolites with the most discriminative pattern of expression that might be proposed as putative biomarkers.

Applying feature importance calculation to the best model obtained on BRAIN dataset, yielded the top four metabolites that correspond to different isotopes and adducts of 2-hydroxyglutarate, a specific product of mutated glial cells, as already found in the previous study (figure 4A) [17]. Mutations of isocitrate dehydrogenase (IDH) enzyme can produce high levels of 2-hydroxyglutarate to inhibit glioma stem cell differentiation, increase tumor microenvironment formation and produce high levels of hypoxia-inducible factor-1α to promote glioma invasion. Mutations in the IDH enzyme worsen the prognosis of gliomas. It is therefore important to distinguish between the two types of glial tumor in order to tailor treatments and improve prognosis [27].

**Figure 4:**
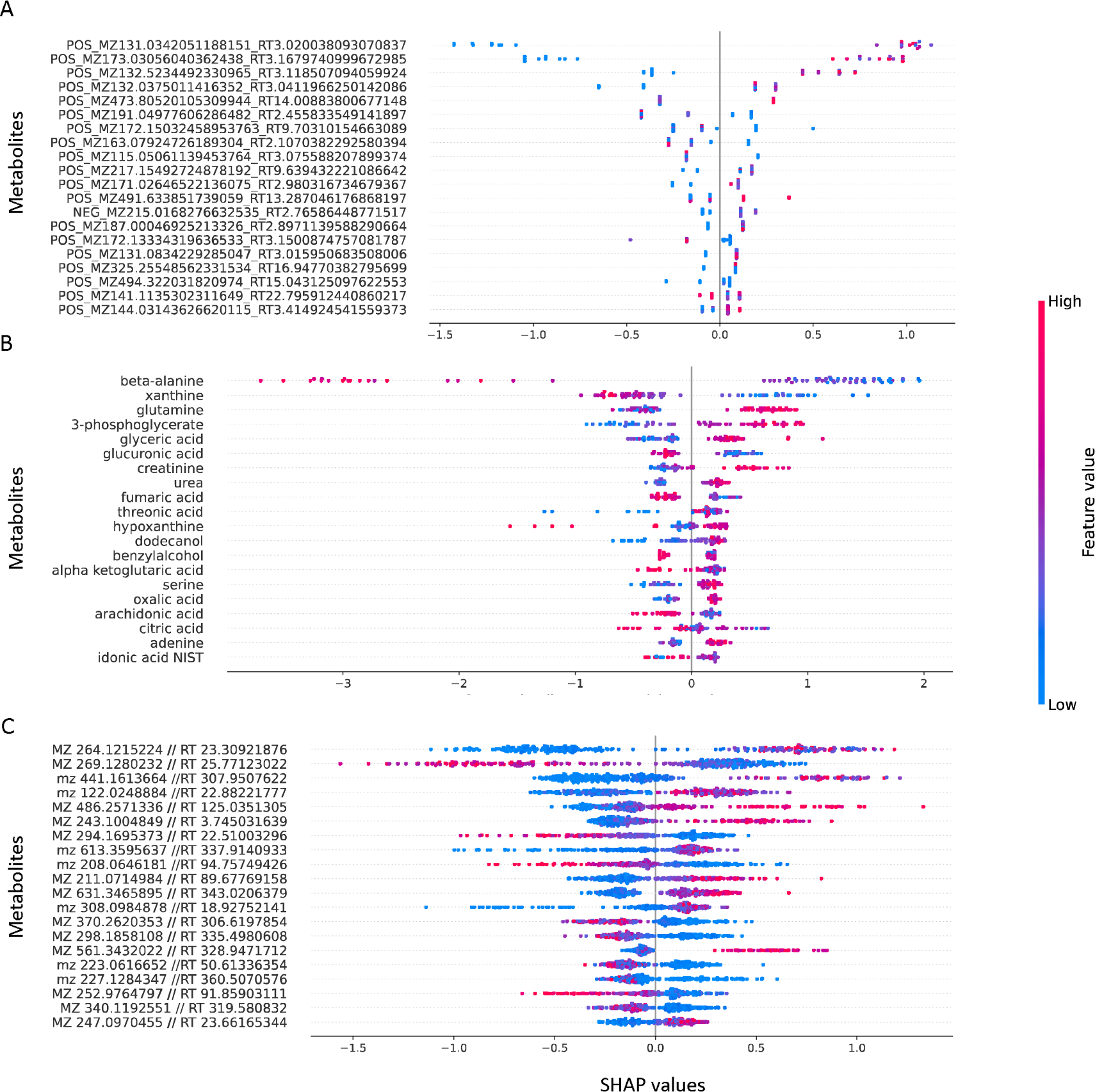
Feature importance bees-plots obtained by using Shapley Additive exPlanations (SHAP). Summary plots are reported for the top contributing features according with SHAP analysis on the more robust model for our study, NFE, namely XGBoost coupled with feature selection for BRAIN dataset (A), BREAST dataset (B) and LUNG dataset (C). On x axis, SHAP values are represented, which is a score representing the contribution of a feature to the model. On y-axis, there are the top metabolites/features that contribute to the model.

For the BREST dataset, the aim is to distinguish the cancer status depending on the hormone receptor (ER) that is crucial for determining which treatment to administer to patients. Indeed, hormone therapy drugs can be used for ER+ breast cancer samples but will be ineffective for ER- breast tumors. The two best performing methods for discriminating ER+ and ER- tumors are HDDA and MPPCA, however they do not allow to calculate feature importance and hence to provide potential biomarkers. Since the model that combines feature selection and XGBoost classifier achieved similar performances as MPPCA and HDDA models, with an average balanced accuracy of 87.2% (±2.8%), an average recall of 89.6% (±2.9%), an average specificity of 66.4% (±10.9%) and an average F1 score of 92.5% (±0.6%) (figure 2B), we calculated the feature importance for this configuration. The top two most contributing metabolites are beta-alanine and the xhantine (figure 4B). Higher values of both metabolites seem to have negative impact on the model and lower values have a positive impact on the model. This means that high concentrations of both metabolites lead to lower risks of developing ER+ tumors compared with low concentrations that yields to high risks of developing ER+ tumors. Both metabolites have already been shown to have significantly different concentrations in ER+ and ER- breast tumors and they have already been suggested to be used as biomarkers to distinguish the two types of breast tumors. Furthermore, the glutamine, which is the third metabolite in feature importance, is derived from glutamic acid and increased concentrations of glutamic acid indicate higher glutaminolysis, a key feature of metabolic changes in cancer cells [20].

Then we inspected the contribution of the metabolites to the best classification model for the LUNG dataset. Identify potential biomarkers for this cancer is fundamental since early detection is pivotal for the treatment of this aggressive cancer. The metabolite that ranked at the first position in feature importance corresponds to the creatine riboside (figure 4C). This metabolite was described as the most important metabolite to discriminate between lung cancer patients and healthy individuals [22].

Overall, SHAP analysis on the best performing models allowed to identify the most contributing metabolites to discriminate the samples depending on the phenotype and is a valid tool to identify potential biomarkers.

## Discussion and Conclusions

In this paper, we have proposed a workflow to classify two groups of samples using feature selection and feature extraction methods. We demonstrated that cross-validation is the best strategy for achieving good classification performances with unbalanced and small datasets, that are very common in the biomedical field. Interestingly, we obtained better results with the two unbalanced datasets, which had few samples, compared to the LUNG dataset, which had over 1,000 samples and two equally represented classes. The drawback of implementing cross-validation consists of a considerable extension of training time and a substantial computational cost, requiring significant processing power.

We showed that, independently on the classification technique applied, feature selection is a necessary step before performing feature extraction method to improve the performances. On the other hand, feature selection can eliminate important features, therefore the feature selection method must be chosen very carefully and adapted to the data and biomedical model under investigation.

Unexpectedly, we have found that performances of linear and non-linear methods are similar. Our hypothesis is that the metabolites measurements used in this study are not entangled among each other as for other complex diseases or other omics, thus both types of techniques are able to capture the essential characteristics to classify the patients depending on their phenotype. Indeed, urine sample is influenced by many factors as race, age, lifestyle (diet, smoke, physical activity) and microbiota. We might expect that in other cases, performances can be different.

Importantly, metabolomics has the potentiality to be a clinical tool for detecting cancer as early as possible to improve survival rates, and for distinguishing between two types of tumors to tailor treatment and improve efficacy. Therefore, the possibility to use techniques that allow to finding cancer biomarkers using inexpensive, non-invasive methods might be preferred if the performances are similar, as showed in our study.

The integration of metabolomics with other omics approaches, such as transcriptomics and proteomics would offer a global perspective not only in cancer biology but in any complex disease, revealing metabolic dysregulations and their interaction with other molecular pathways. In this scenario the datasets will be even more unbalanced than using a single omics because the number of features will increase dramatically, reaching several thousand depending on the omics, and would be hardly comparable to the number of patients. In such perspective the use of features selection and feature extraction methods will become indispensable, and we believe that the guidelines set on this study would help to benchmark these techniques on more complex datasets paving the way toward a more effective precision medicine using multi-omics data.

## Author contributions

JL: Data curation; Formal analysis; Methodology; Software; Visualization; Roles/Writing - original draft; ENF: Data curation; Formal analysis; Methodology; Software; Writing - review & editing; SB: Conceptualization; Supervision; Roles/Writing - original draft.

