## supplementary materials for "Benchmarking feature selection and feature extraction methods to improve the performances of machine-learning algorithms for patient classification using metabolomics biomedical data"

Supplementary Table 1: Classification results from feature extraction methods for BRAIN dataset.

In the table, the mean and standard deviation is reported for each feature extraction method with and without feature selection for BRAIN dataset. Abbreviations for feature extraction method are: NFE: No Feature Extraction; HDDA: High dimensional discriminant analysis; MPPCA: Mixture of Probalistic PCA; PCA: Principal Component Analysis (PCA); KPCA: Kernel PCA; GPLVM: Gaussian Process Latent Variable Modeling.

| **Feature extraction method** | **Classifier** | **Feature selection** | **Balanced Accuracy** | **Precision** | **Recall** | **F1 Score** | **ROC AUC** | **Specificity** |
| --- | --- | --- | --- | --- | --- | --- | --- | --- |
| NFE | Xgboost | No | 89.4 (±6.4) | 90.7 (±5.2) | 92.0 (±7.4) | 89.9 (±6.6) | 95.1 (±4.4) | 88.9 (±10.8) |
| NFE | Xgboost | Yes | 89.6 (±6.9) | 91.0 (±5.3) | 92.1 (±7.8) | 90.4 (±6.5) | 94.6 (±4.5) | 88.9 (±12.0) |
| HDDA | Statistical | No | 50.7 (±3.9) | 57.6 (±5.2) | 70.1 (±6.5) | 62.9 (±3.2) | 52.4 (±5.1) | 31.4 (±13.7) |
| HDDA | Statistical | Yes | 82.7 (±7.9) | 82.1 (±6.3) | 91.6 (±9.8) | 86.5 (±7.4) | 87.1 (±12.9) | 73.9 (±9.3) |
| MPPCA | Statistical | No | 58.3 (±9.8) | 63.0 (±8.6) | 71.7 (±11.0) | 67.0 (±9.4) | 59.4 (±7.7) | 45.0 (±11.0) |
| MPPCA | Statistical | Yes | 75.8 (±15.3) | 75.5 (±12.9) | 87.8 (±15.0) | 81.1 (±13.8) | 82.8 (±17.5) | 63.9 (±16.0) |
| PCA | Xgboost | No | 67.4 (±9.1) | 74.7 (±8.0) | 69.8 (±7.7) | 81.3 (±11.8) | 74.9 (±9.9) | 53.4 (±13.8) |
| KPCA | Xgboost | No | 53.9 (±11.9) | 59.5 (±12.7) | 59.9 (±11.2) | 60.3 (±16.5) | 54.5 (±12.7) | 47.6 (±15.7) |
| GPLVM | Statistical | No | 47.3 (±7.9) | 54.6 (±6.5) | 84.3 (±20.7) | 65.9 (±11.4) | 57.9 (±6.1) | 10.3 (±8.2) |
| GPLVM | Statistical | Yes | 87.7 (±8.4) | 92.0 (±5.8) | 86.2 (±16.9) | 88.3 (±10.4) | 93.7 (±7.8) | 89.2 (±9.1) |

Supplementary Table 2: Classification results from feature extraction methods for BREAST dataset.

In the table, the mean and standard deviation is reported for each feature extraction method with and without feature selection for BREAST dataset. Abbreviations for feature extraction method are: NFE: No Feature Extraction; HDDA: High dimensional discriminant analysis; MPPCA: Mixture of Probalistic PCA; PCA: Principal Component Analysis (PCA); KPCA: Kernel PCA; GPLVM: Gaussian Process Latent Variable Modeling

Two FA were performed, the first with 7 features and the second with 10 features.

| **Feature extraction method** | **Classifier** | **Feature selection** | **Balanced Accuracy** | **Precision** | **Recall** | **F1 Score** | **ROC AUC** | **Specificity** |
| --- | --- | --- | --- | --- | --- | --- | --- | --- |
| NFE | Xgboost | No | 79.2 (±5.0) | 91.4 (±2.0) | 89.1 (±2.7) | 93.8 (±2.9) | 87.1 (±5.3) | 64.7 (±9.7) |
| NFE | Xgboost | Yes | 80.2 (±5.2) | 91.7 (±1.8) | 89.6 (±2.9) | 94.0 (±2.7) | 88.2 (±3.7) | 66.4 (±10.9) |
| HDDA | Statistical | No | 79.0 (±3.7) | 89.7 (±2.2) | 89.2 (±2.5) | 89.4 (±1.7) | 86.0 (±3.2) | 68.8 (±6.9) |
| HDDA | Statistical | Yes | 84.0 (±6.0) | 92.0 (±4.2) | 93.1 (±3.4) | 92.5 (±0.6) | 90.7 (±3.3) | 74.8 (±15.3) |
| MPPCA | Statistical | No | 77.8 (±4.4) | 88.9 (±2.6) | 89.7 (±3.3) | 89.3 (±2.1) | 86.6 (±3.7) | 65.8 (±8.3) |
| MPPCA | Statistical | Yes | 84.2 (±4.5) | 92.6 (±2.9) | 90.7 (±4.9) | 91.6 (±2.5) | 89.9 (±4.1) | 77.8 (±9.8) |
| PCA | Xgboost | No | 74.5 (±4.6) | 90.4 (±2.1) | 86.4 (±2.2) | 94.8 (±3.7) | 84.6 (±3.3) | 54.2 (±9.4) |
| FA1 - 7 features | Xgboost | Yes | 78.0 (±4.4) | 90.8 (±1.8) | 88.5 (±2.4) | 93.3 (±3.3) | 87.2 (±3.5) | 62.6 (±9.6) |
| FA2 - 10 features | Xgboost | Yes | 73.7 (±5.0) | 89.3 (±1.9) | 86.3 (±2.5) | 92.6 (±3.0) | 84.7 (±3.6) | 54.8 (±10.0) |
| KPCA | Xgboost | No | 57.5 (±5.7) | 80.6 (±3.3) | 78.9 (±2.7) | 82.7 (±6.2) | 65.9 (±6.5) | 32.2 (±12.3) |
| GPLVM | Statistical | No | 58.2 (±3.9) | 78.5 (±1.4) | 100 (±0) | 87.9 (±0.9) | 88.2 (±3.3) | 16.5 (±5.8) |
| GPLVM | Statistical | Yes | 71.6 (±4.8) | 84.5 (±2.3) | 98.5 (±1.9) | 91.0 (±1.2) | 90.0 (±2.8) | 44.7 (±10.7) |

Supplementary Table 3: Classification results from feature extraction methods for LUNG dataset.

In the table, the mean and standard deviation is reported for each feature extraction method with and without feature selection for LUNG dataset. Abbreviations for feature extraction method are: NFE: No Feature Extraction; HDDA: High dimensional discriminant analysis; MPPCA: Mixture of Probalistic PCA; PCA: Principal Component Analysis (PCA); KPCA: Kernel PCA; GPLVM: Gaussian Process Latent Variable Modeling

Two FA were performed, the first with 17 features and the second with 44 features.

| **Feature extraction method** | **Classifier** | **Feature selection** | **Accuracy** | **Precision** | **Recall** | **F1 Score** | **ROC AUC** | **Specificity** |
| --- | --- | --- | --- | --- | --- | --- | --- | --- |
| NFE | Xgboost | No | 78.0 (±2.2) | 75.2 (±2.4) | 79.6 (±3.1) | 71.3 (±2.6) | 77.6 (±2.2) | 83.9 (±2.9) |
| NFE | Xgboost | Yes | 78.4 (±2.4) | 76.2 (±2.7) | 78.7 (±2.9) | 73.8 (±3.1) | 78.2 (±2.4) | 82.5 (±2.6) |
| HDDA | Statistical | No | 63.3 (±2.7) | 59.4 (±3.8) | 69.7 (±13.5) | 63.3 (±5.0) | 68.5 (±3.3) | 57.5 (±13.9) |
| HDDA | Statistical | Yes | 64.9 (±3.1) | 60.6 (±4.8) | 74.0 (±9.7) | 66.0 (±3.1) | 72.6 (±2.5) | 56.8 (±12.1) |
| MPPCA | Statistical | No | 64.6 (±2.3) | 62.3 (±3.8) | 61.1 (±4.2) | 61.5 (±2.6) | 67.6 (±2.1) | 67.7 (±4.5) |
| MPPCA | Statistical | Yes | 68.5 (±2.0) | 68.3 (±2.4) | 60.4 (±6.7) | 63.9 (±3.8) | 75.1 (±2.4) | 75.6 (±3.7) |
| PCA | Xgboost | No | 67.6 (±2.3) | 62.2 (±3.1) | 68.3 (±3.0) | 57.1 (±3.9) | 73.8 (±2.1) | 76.7 (±3.0) |
| FA1 - 17 features | Xgboost | Yes | 73.4 (±2.2) | 70.1 (±3.3) | 73.9 (±3.0) | 67.0 (±5.8) | 80.9 (±2.6) | 79.1 (±4.0) |
| FA2 - 44 features | Xgboost | Yes | 72.7 (±2.2) | 69.4 (±3.1) | 72.7 (±2.3) | 66.6 (±5.2) | 80.2 (±2.1) | 78.0 (±3.1) |
| KPCA | Xgboost | No | 64.3 (±2.1) | 59.7 (±2.4) | 63.1 (±2.7) | 56.7 (±3.0) | 69.3 (±2.2) | 70.9 (±3.2) |
| GPLVM | Statistical | No | 72.4 (±2.7) | 75.6 (±4.3) | 59.8 (±5.6) | 66.6 (±3.5) | 80.3 (±2.6) | 83.4 (±3.4) |
| GPLVM | Statistical | Yes | 75.6 (±2.2) | 76.6 (±4.2) | 68.4 (±4.6) | 72.1 (±2.5) | 83.6 (±1.8) | 82.0 (±3.8) |
